## Supplementary information for "Synthetic genotype networks"

**Supplementary Table 1.** Plasmids used in this study. †Plasmids from Santos-Moreno & Schaerli 2019, ACS Synth Biol. 8, 1691-1697. *Plasmids from Santos-Moreno et al. 2020, Nat. Commun. 11, 2746. The rest of the plasmids were constructed in this study.

| Name | Resistance | Ori | Phenotype / features | Usage |
|---|---|---|---|---|
| pC-0 | Kanamycin | ColA | Empty multiple cloning site | Used for non-fluorescent control bacteria† |
| pJ1996_v2 | Spectinomycin | CloDF13 | dCas9 & Csy4 | Used to express dCas9 and csy4 required for all GRNs† |
| 4.1 | Kanamycin | ColA | BLUE-stripe | Fig. 3 |
| 3.3 | Kanamycin | ColA | BLUE-stripe | Fig. 3 |
| 3.2 | Kanamycin | ColA | BLUE-stripe | Fig. 3 |
| 3.1 | Kanamycin | ColA | BLUE-stripe | Fig. 3 |
| 2-OS.1 | Kanamycin | ColA | Oscillator | Fig. 8 |
| 2c-P.1 | Kanamycin | ColA | GREEN-stripe | Fig. 5 |
| 2c-NF.1 | Kanamycin | ColA | Non-functional | Fig. 7 |
| 2c-CT.6 | Kanamycin | ColA | GREEN-stripe | Fig. S1 |
| 2c-CT.5 | Kanamycin | ColA | GREEN-stripe | Fig. S1 |
| 2c-CT.4 | Kanamycin | ColA | GREEN-stripe | Fig. S1 |
| 2c-CT.3 | Kanamycin | ColA | GREEN-stripe | Fig. S1 |
| 2c-CT.2 | Kanamycin | ColA | BLUE-stripe | Fig. S1 |
| 2c-CT.1 | Kanamycin | ColA | BLUE-stripe | Fig. S1 |
| 2c.8 | Kanamycin | ColA | BLUE-stripe | Fig. 3 |
| 2c.7 | Kanamycin | ColA | BLUE-stripe | Fig. 3 |
| 2c.6 | Kanamycin | ColA | BLUE-stripe | Fig. 3 |
| 2c.5 | Kanamycin | ColA | BLUE-stripe | Fig. 3 |
| 2c.4 | Kanamycin | ColA | BLUE-stripe | Fig. 3 |
| 2c.3 | Kanamycin | ColA | BLUE-stripe | Fig. 3 |
| 2c.2 | Kanamycin | ColA | BLUE-stripe | Fig. 3 |
| 2c.1 | Kanamycin | ColA | BLUE-stripe | Fig. 3 |
| 2b.2 | Kanamycin | ColA | GREEN-stripe | Fig. 2 |
| 2b.1 | Kanamycin | ColA | GREEN-stripe | Fig. 2 |
| 2a.1 | Kanamycin | ColA | GREEN-stripe | Fig. 2 |
| 1-OS.3 | Kanamycin | ColA | Oscillator | Fig. 8* |
| 1-OS.2 | Kanamycin | ColA | Oscillator | Fig. 8 |
| 1-OS.1 | Kanamycin | ColA | Oscillator | Fig. 8 |
| 1.4 | Kanamycin | ColA | GREEN-stripe | Fig. 2 |
| 1.3 | Kanamycin | ColA | GREEN-stripe | Fig. 2 |
| 1.2 | Kanamycin | ColA | GREEN-stripe | Fig. 2 |
| 1.1 | Kanamycin | ColA | GREEN-stripe | Fig. 2* |

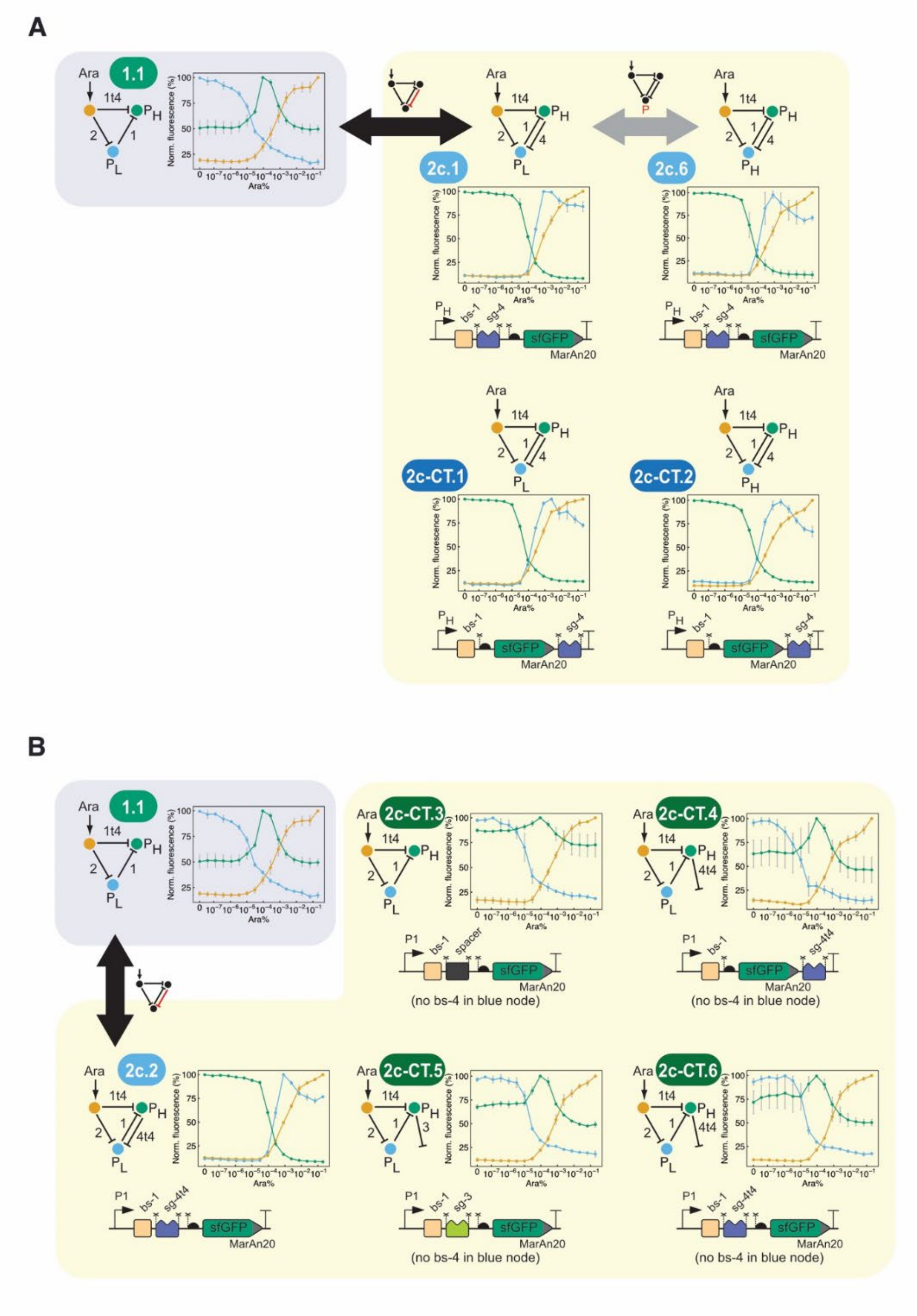

**Supplementary Fig. 1.** Controls supporting the GREEN-stripe to BLUE-stripe transition of topology 2c (related to **Fig. 2** and **3**). **A**. GREEN-to-BLUE transition is not due to polar effects derived from the presence of a new sgRNA upstream sfGFP. Designs 2c-CT.1 and 2c-CT.2 are equivalent to 2c.1 and 2c.6 but with the fourth sgRNA (sgRNA-4) downstream sfGFP and yet produce a BLUE-stripe phenotype. **B**. GREEN-to-BLUE transition requires a functional repression from the green node to the blue one. Whenever the repression is disrupted (designs 2c-CT.3, 2c-CT.4, 2c-CT.5 and 2c-CT.6), the resulting phenotype is that of a GREEN-stripe. Data from three biological replicates. Symbols as in **Fig. 1** and **2**.

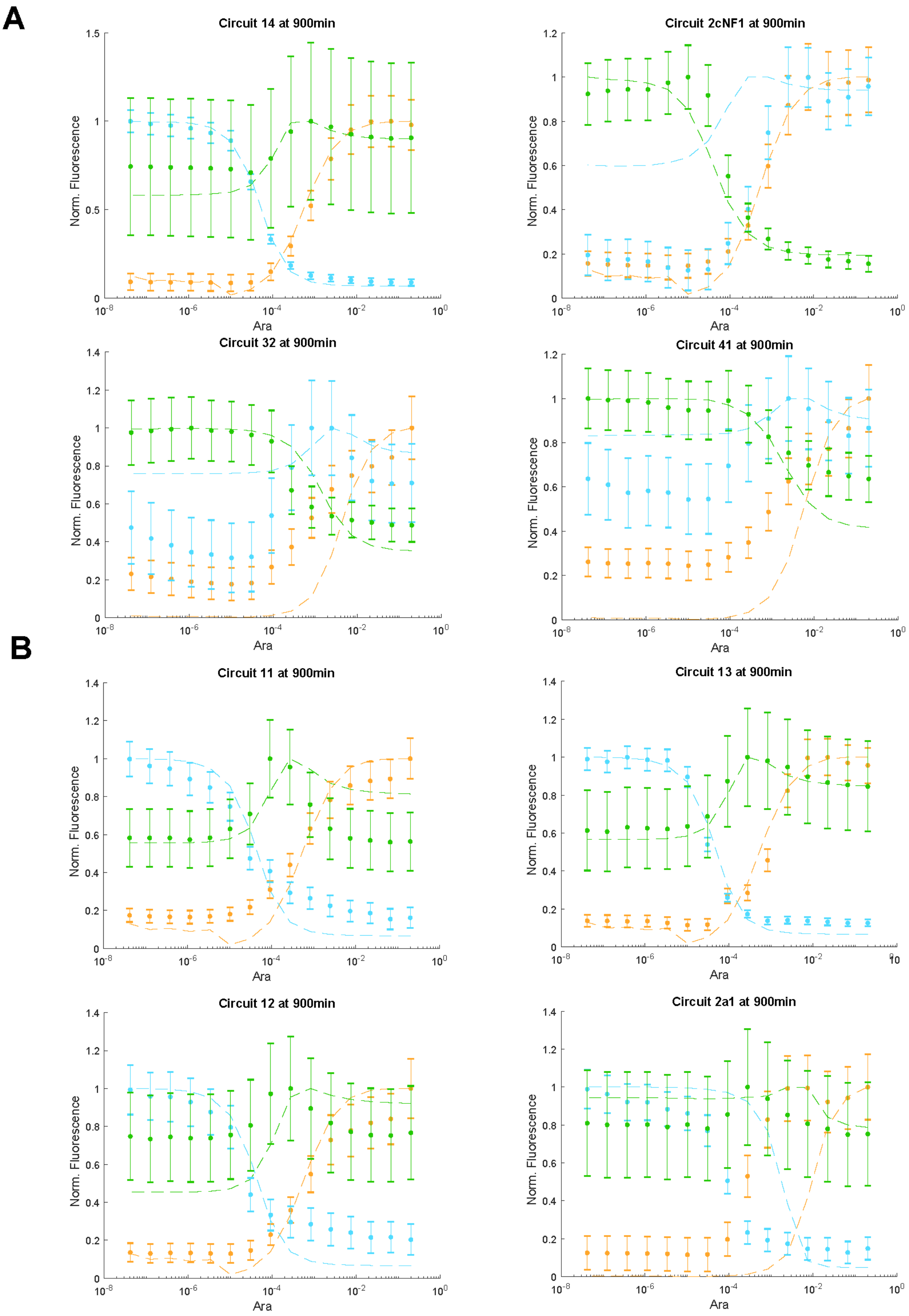
A
Circuit 14 at 900min
Circuit 2cNF1 at 900min
Circuit 32 at 900min
Circuit 41 at 900min
B
Circuit 11 at 900min
Circuit 13 at 900min
Circuit 12 at 900min
Circuit 2a1 at 900min
Norm. Fluorescence
Ara

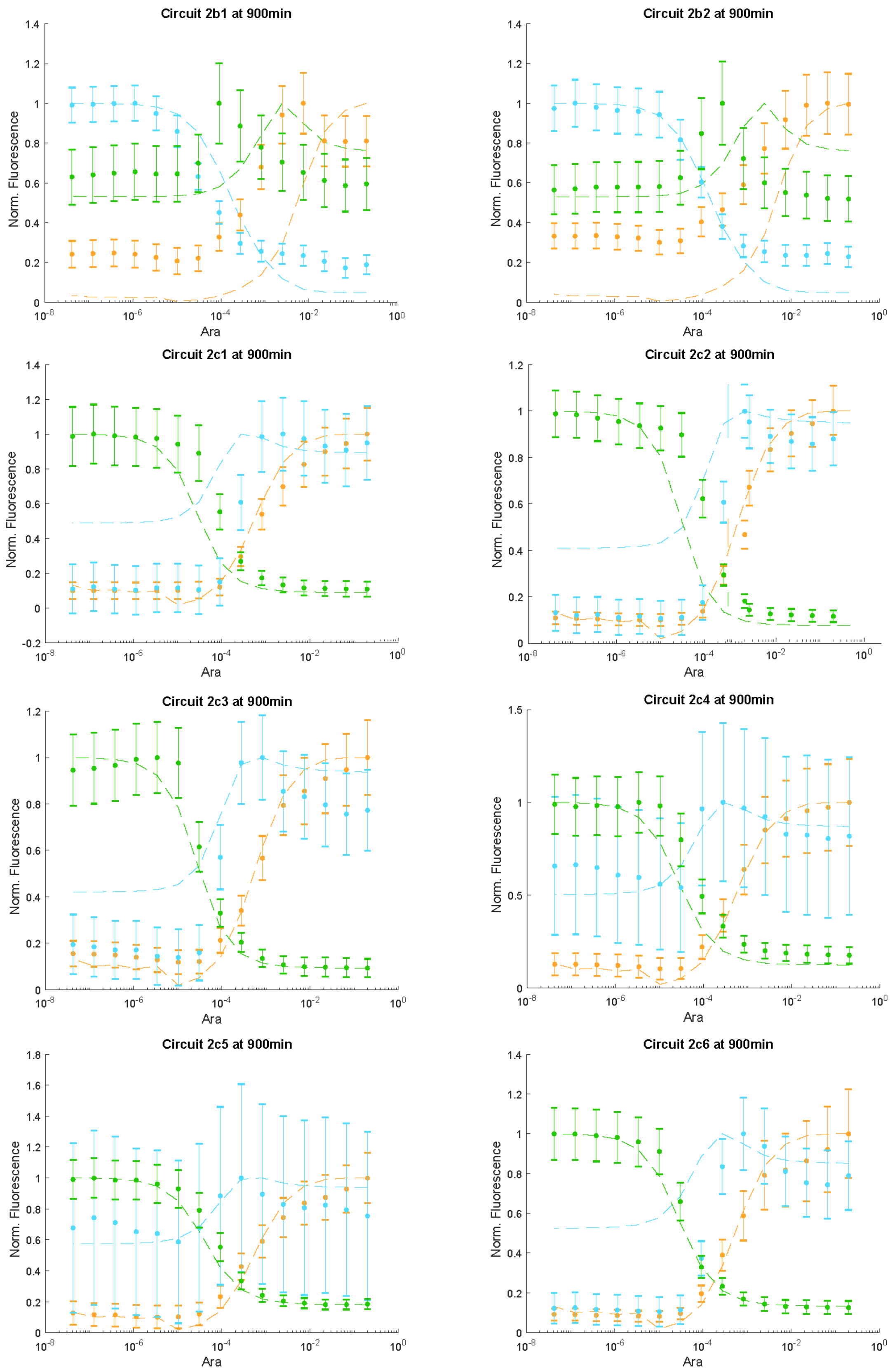
Circuit 2b1 at 900min
Circuit 2b2 at 900min
Circuit 2c1 at 900min
Circuit 2c2 at 900min
Circuit 2c3 at 900min
Circuit 2c4 at 900min
Circuit 2c5 at 900min
Circuit 2c6 at 900min
Norm. Fluorescence
Ara

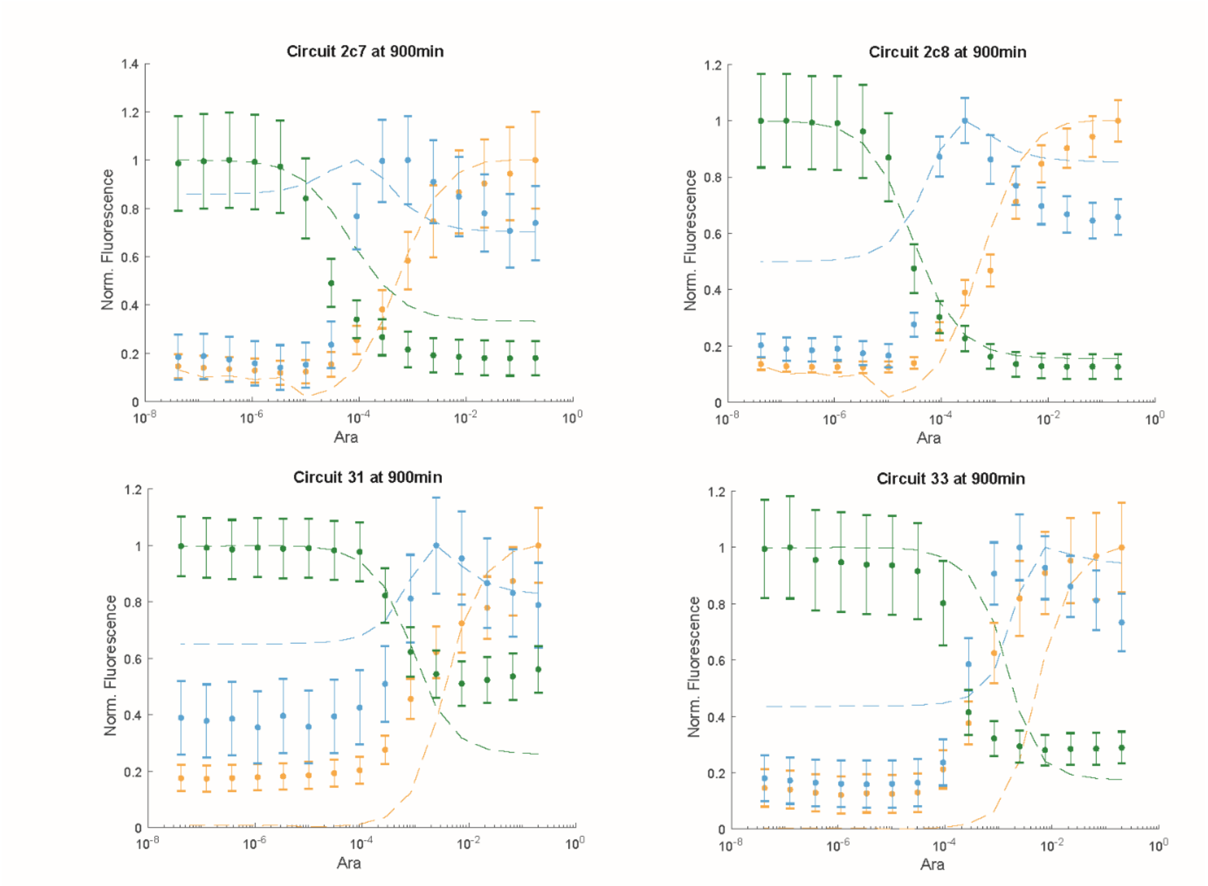


**Supplementary Fig. 2.** Comparing models with data for stripe generation for different arabinose inducer concentrations at 900min. **A**. Experimental data (symbols) and simulation results (lines) for GRNs used in the parameter estimation process. Colors indicate node A (orange), B (blue), and C (green). Error bars show s.d. based on error propagation. **B**. As **A.**, for GRNs used in model validation.

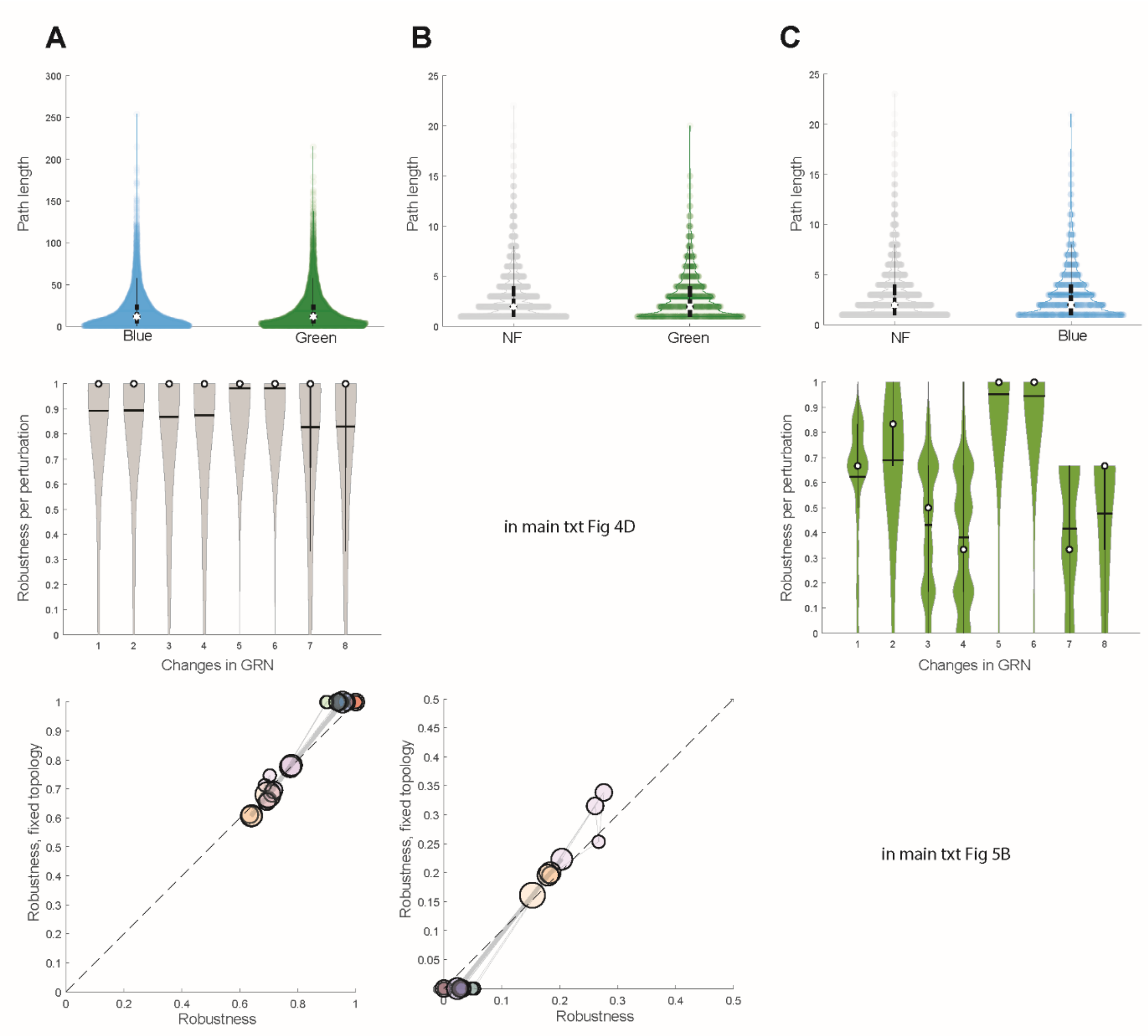


**Supplementary Fig. 3.** Genotype network characteristics for the three stripe phenotypes. **A.** Characteristics for the NF phenotype. We show the path length for obtaining a BLUE-stripe (2) or a GREEN-stripe (3) when starting with a NF phenotype (**top**). Robustness per perturbation (**middle**) is computed for changes in a randomly sampled set of NF GRNs (n = 2000). **Bottom**: robustness for the NF phenotype per topology over all neighboring genotypes vs only genotypes with a fixed topology. **B.** and **C.** Corresponding results for the BLUE- and GREEN-stripe GRNs, respectively.

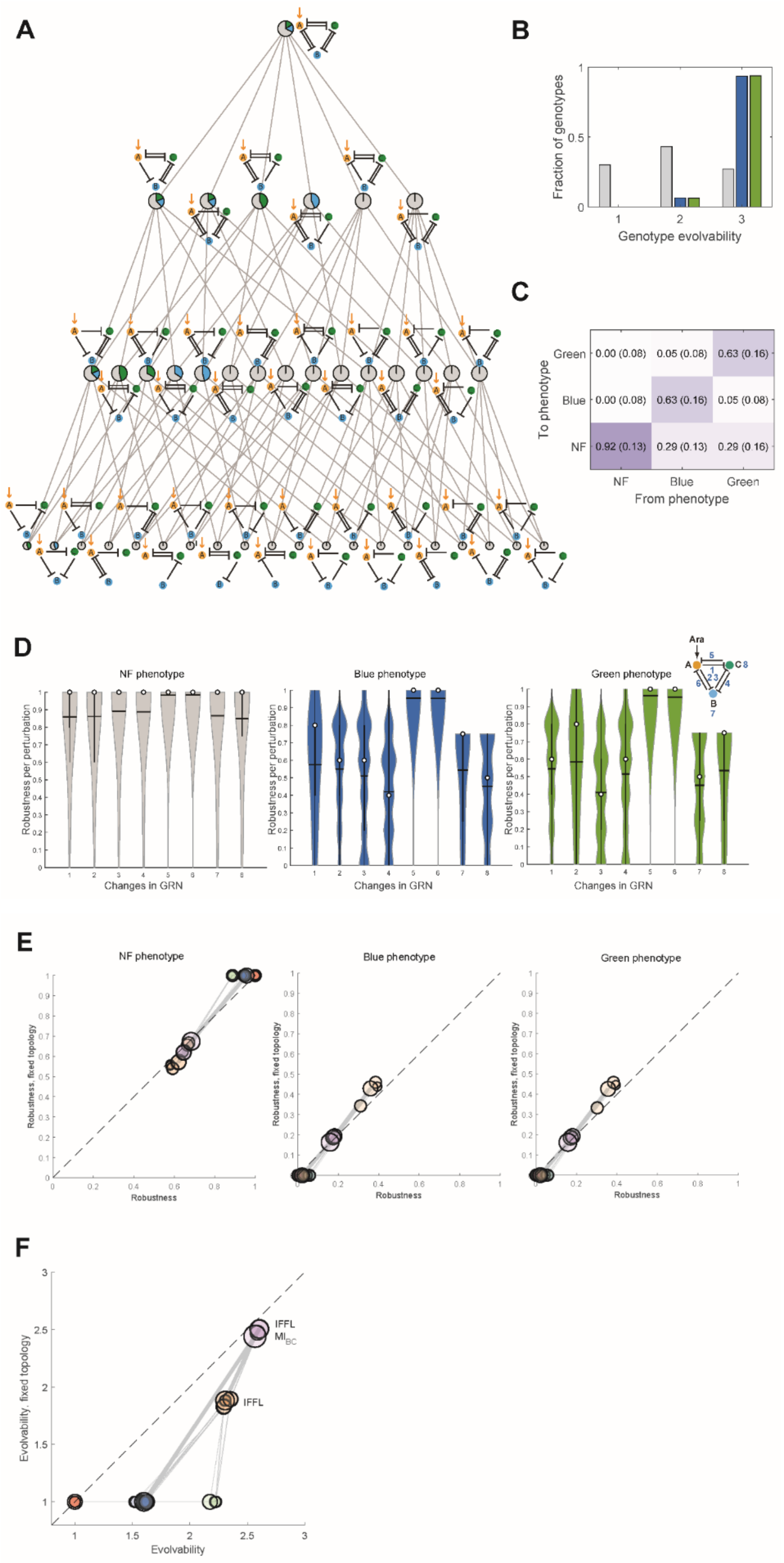


**Supplementary Fig. 4.** Analysis results for artificial inhibition and promoter strengths. **A**. Network topologies with their neighbor relations (similar to **Fig. 4A**). **B**. Distributions of evolvability for genotypes with BLUE-, GREEN- and NF-stripe (similar to **Fig. 4B**). **C**. Transition frequencies between indicated phenotypes (similar to **Fig. 4C**). **D**. Robustness per perturbation for NF-, BLUE-, and GREEN-stripe (similar to **Fig. 4E** and **SF3**). **E**. Robustness for the three phenotypes per topology over all neighboring genotypes vs only genotypes with a fixed topology (similar to **Fig. 5B** and **SF3**). **F**. Average predicted phenotypic evolvability per topology over all neighboring genotypes vs only genotypes with a fixed topology (similar to **Fig. 5A**).

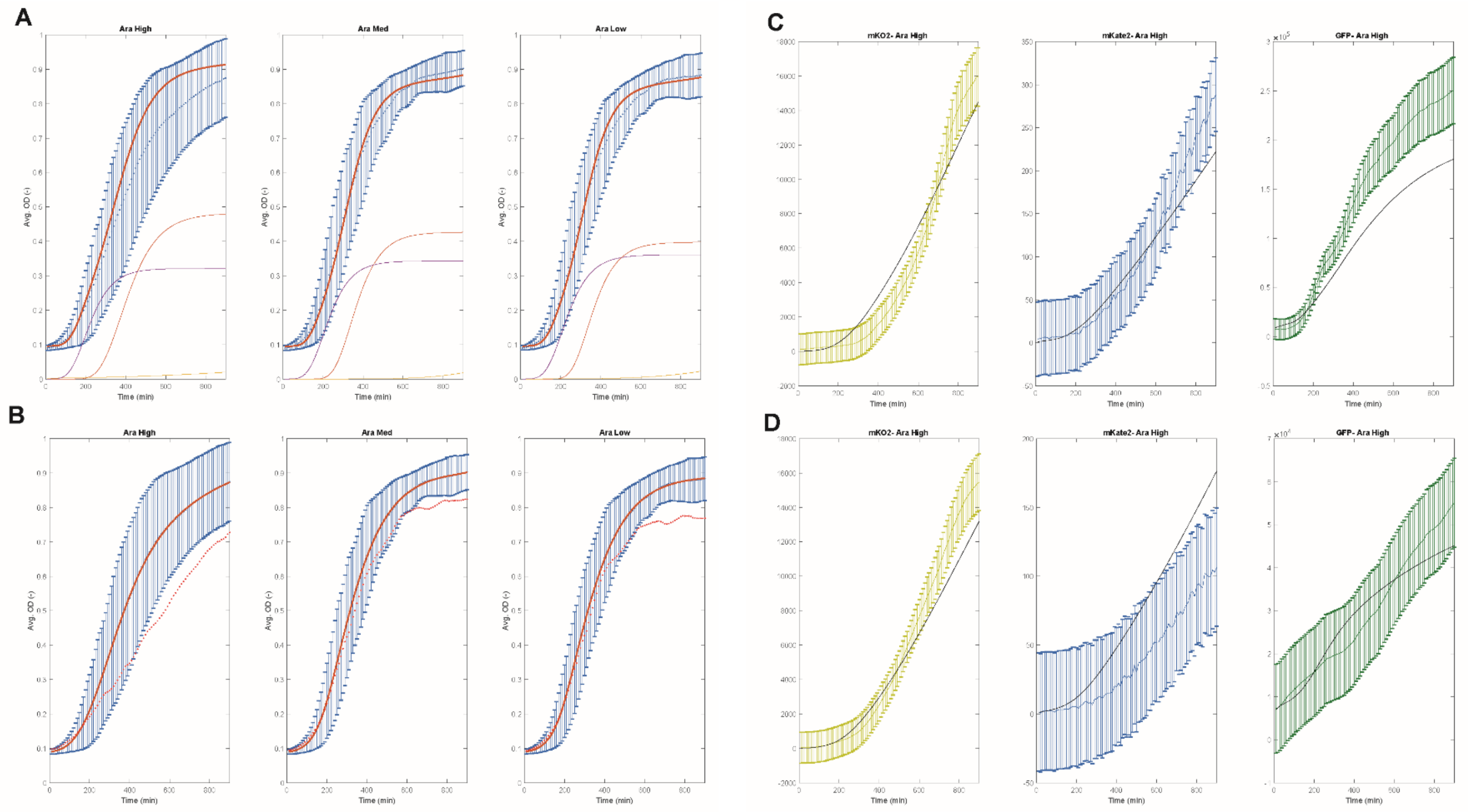
A
Ara High
Ara Med
Ara Low
Avg. OD (-)
Time (min)
B
Ara High
Ara Med
Ara Low
Avg. OD (-)
Time (min)
C
mKO2- Ara High
mKate2- Ara High
GFP- Ara High
Time (min)
D
mKO2- Ara High
mKate2- Ara High
GFP- Ara High
Time (min)

**E**

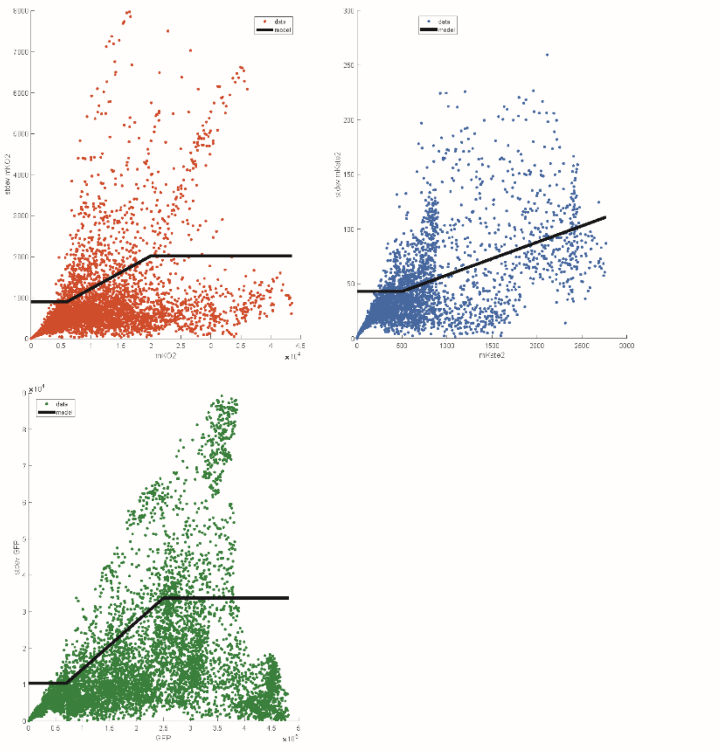

**Supplementary Fig. 5.** Analysis examples of experimental data with models. **A.** Growth model (orange) and OD data (blue, mean ± s.d. over 3 independent experiments) between 0min and 900min after induction with arabinose with points every 10min for GRN 4.1 (used in the estimation procedure) during high, intermediate and zero arabinose induction. The growth model is a superposition of three sigmoidal functions (additional colored lines). **B.** The averaged OD dataset (blue) used for the GRNs that have four inhibitions with the estimated growth model (orange). The red line corresponds to the GRN 2c.4 OD similar to A. **C.** ODE model trajectories (solid black line) with the relevant experimental data for the three reporters (yellow: mKO2, blue: mKate2, green: GFP; mean ± s.d., where error bars represent the measurement model used) for high arabinose induction during 0min to 900min (with points every 10min) for  GRN4.1. **D.** Predicted ODE model trajectories for GRN 2c4 using the average growth model compared to the available experimental data at high arabinose. The axis and color-coding is the same as in C. **E.** Estimation of measurement variances for the three reporters used. For each reporter, the fluorescence data from all experiments was pooled, binned my mean fluorescence, and the standard deviation was plotted against mean fluorescence of the binned data (color-coded circles). Piecewise linear models (black lines) were used per reporter.

**Supplementary Table 2:** Topologies with their stripe phenotype prevalence (from **Fig. 4A**) and the clustering ID (from **Fig. 5A**,**B**). Each topology is represented by the presence (1) or absence (0) of an inhibition, without specifying the experimental sgRNA or promoter used. The indexing of each 0,1 is similar to **Fig. 4E**, but here without the last two elements for the promoter efficiencies. The NF, BLUE, and GREEN columns show the percentage of each stripe per topology. In the column clusterID, the numbers 1 to 5 refer to the colors red, blue, green, purple and orange, respectively in **Fig. 5A**,**B**.

| Topology | NF | BLUE | GREEN | clusterID |
|---|---|---|---|---|
| 0 0 0 1 1 1 | 1 | 0 | 0 | 1 |
| 0 0 1 0 1 1 | 1 | 0 | 0 | 1 |
| 0 0 1 1 0 1 | 1 | 0 | 0 | 1 |
| 0 0 1 1 1 0 | 1 | 0 | 0 | 1 |
| 0 0 1 1 1 1 | 1 | 0 | 0 | 1 |
| 0 1 0 0 1 1 | 1 | 0 | 0 | 1 |
| 0 1 0 1 0 1 | 1 | 0 | 0 | 2 |
| 0 1 0 1 1 0 | 1 | 0 | 0 | 2 |
| 0 1 0 1 1 1 | 1 | 0 | 0 | 2 |
| 0 1 1 0 0 1 | 1 | 0 | 0 | 2 |
| 0 1 1 0 1 0 | 1 | 0 | 0 | 2 |
| 0 1 1 0 1 1 | 1 | 0 | 0 | 2 |
| 0 1 1 1 0 0 | 1 | 0 | 0 | 2 |
| 0 1 1 1 0 1 | 1 | 0 | 0 | 2 |
| 0 1 1 1 1 0 | 1 | 0 | 0 | 2 |
| 0 1 1 1 1 1 | 1 | 0 | 0 | 2 |
| 1 0 0 0 1 1 | 1 | 0 | 0 | 1 |
| 1 0 0 1 0 1 | 1 | 0 | 0 | 2 |
| 1 0 0 1 1 0 | 1 | 0 | 0 | 2 |
| 1 0 0 1 1 1 | 1 | 0 | 0 | 2 |
| 1 0 1 0 0 1 | 1 | 0 | 0 | 2 |
| 1 0 1 0 1 0 | 1 | 0 | 0 | 2 |
| 1 0 1 0 1 1 | 1 | 0 | 0 | 2 |
| 1 0 1 1 0 0 | 1 | 0 | 0 | 2 |
| 1 0 1 1 0 1 | 1 | 0 | 0 | 2 |
| 1 0 1 1 1 0 | 1 | 0 | 0 | 2 |
| 1 0 1 1 1 1 | 1 | 0 | 0 | 2 |
| 1 1 0 0 0 1 | 1 | 0 | 0 | 3 |
| 1 1 0 0 1 0 | 1 | 0 | 0 | 3 |
| 1 1 0 0 1 1 | 1 | 0 | 0 | 3 |
| 1 1 0 1 0 0 | 0.75 | 0.25 | 0 | 4 |
| 1 1 0 1 0 1 | 0.66 | 0.34 | 0 | 4 |
| 1 1 0 1 1 0 | 0.68 | 0.32 | 0 | 4 |
| 1 1 0 1 1 1 | 0.78 | 0.22 | 0 | 4 |
| 1 1 1 0 0 0 | 0.71 | 0 | 0.29 | 4 |
| 1 1 1 0 0 1 | 0.7 | 0 | 0.3 | 4 |
| 1 1 1 0 1 0 | 0.66 | 0 | 0.34 | 4 |
| 1 1 1 0 1 1 | 0.78 | 0 | 0.22 | 4 |
| 1 1 1 1 0 0 | 0.61 | 0.2 | 0.19 | 5 |
| 1 1 1 1 0 1 | 0.61 | 0.2 | 0.19 | 5 |
| 1 1 1 1 1 0 | 0.61 | 0.2 | 0.2 | 5 |
| 1 1 1 1 1 1 | 0.68 | 0.16 | 0.16 | 5 |

**Supplementary Text**

By using the assumptions on the sgRNA and reporter dynamics reported in the Methods section, we can construct all the forty-two different topologies shown in **Fig. 4** of the main text. Below, we provide example models and present more details on the structure and implementation. The notation for the kinetic constants used in these examples is the same as in the Methods section, with the difference now that i and j refer to specific parametrizations.

A 2-node topology model

Here we present the ODE system with the algebraic functions for the 2-node model, assuming that we have:

- One inhibition via expression of $sgRNA_1$ and the corresponding dCas complexes.
- The expression of reporter mKO2 (mRNA: $mRNA_A$, immature/ non-fluorescent protein: $PI_A$, fluorescent protein: $P_A$) and the expression of reporter GFP (mRNA: $mRNA_C$, immature/ non-fluorescent protein: $PI_C$, fluorescent protein: $P_C$).

ODE system

sgRNA dynamics

$$\frac{d[sgRNA_1]}{dt} = f_{\left(Ara_{sgRNA_1}\right)} + b_{sgRNA_1} - d_{RNA}\cdot[sgRNA_1] - k_{f_{ds}}\cdot[dCas]\cdot[sgRNA_1] + k_{r_{ds}}\cdot[dCas:sgRNA_1]$$

$$\frac{d[dCas:sgRNA_1]}{dt} = k_{f_{ds}}\cdot[dCas]\cdot[sgRNA_1] + k_{r_{dsd1}}\cdot[dCas:sgRNA_1:DNA_C]$$
$$-m\cdot[dCas:sgRNA_1] - k_{r_{ds}}\cdot[dCas:sgRNA_1] - k_{f_{dsd1}}\cdot[dCas:sgRNA_1]\cdot[DNA_C]$$

$$\frac{d[dCas:sgRNA_1:DNA_C]}{dt} = k_{f_{dsd1}}\cdot[dCas:sgRNA_1]\cdot[DNA_C] - k_{r_{dsd1}}\cdot[dCas:sgRNA_1:DNA_C]$$
$$-m\cdot[dCas:sgRNA_1:DNA_C]$$

Reporter protein dynamics

$$\frac{d[mRNA_A]}{dt} = f_{\left(Ara_{mRNA_A}\right)} + b_{mRNA_A} - d_{RNA}\cdot[mRNA_A]$$

$$\frac{d[mRNA_C]}{dt} = k_{mRNA_C}\cdot[DNA_C] - d_{RNA}\cdot[mRNA_C]$$

$$\frac{d[PI_A]}{dt} = k_{PI_A} \cdot [mRNA_A] - m_{PI_A} \cdot [PI_A] - (m + d_{PI_A}) \cdot [PI_A]$$

$$\frac{d[P_A]}{dt} = m_{PI_A} \cdot [PI_A] - (m + d_{PI_A}) \cdot [P_A]$$

$$\frac{d[PI_C]}{dt} = k_{PI_C} \cdot [mRNA_C] - m_{PI_C} \cdot [PI_C] - (m + d_{PI_C}) \cdot [PI_C]$$

$$\frac{d[P_C]}{dt} = m_{PI_C} \cdot [PI_C] - (m + d_{PI_C}) \cdot [P_C]$$

Algebraic equations

$$[dCas] = [dCas_{total}] - [dCas : sgRNA_1] - [dCas : sgRNA_1 : DNA_C]$$

$$[DNA_C] = [DNA_{C_{total}}] - [dCas : sgRNA_1 : DNA_C]$$

$$\mathrm{f}_{(\mathrm{Ara}_{\mathrm{sgRNA}_1})} = \frac{Ara^n}{Ara^n + K_m{}^n} \cdot k_2 \cdot [\mathrm{DNA}_{A_{total}}] \cdot (1 - \mathrm{b}_{\mathrm{sgRNA}_1})$$

$$\mathrm{f}_{(\mathrm{Ara}_{\mathrm{mRNA}_A})} = \frac{Ara^n}{Ara^n + K_m{}^n} \cdot k_2 \cdot [\mathrm{DNA}_{A_{total}}] \cdot (1 - \mathrm{b}_{\mathrm{mRNA}_A})$$

$$G(t) = p_1 + \sum_{k=1}^{3} \frac{p_2^k}{\left(1 + p_3^k e^{-t \cdot p_4^k}\right)^{1/p_5^k}}$$

$$m(t) = \frac{dG(t)}{dt}$$

<u>A 3-node topology model with three inhibitions</u>

Here we present the ODE system with the algebraic functions for the 3-node model, assuming that we have:

- Three inhibitions via expression of $sgRNA_1$, $sgRNA_2$ and $sgRNA_3$ and the corresponding dCas complexes. Arabinose induces the expression of $sgRNA_1$ and $sgRNA_2$, which inhibit the expression of the constitutively expressed green and blue node, respectively. The blue node induces the expression of $sgRNA_3$, which in turn inhibits the green node.
- The expression of reporter mKO2 (mRNA: $mRNA_A$, immature/ non-fluorescent protein: $PI_A$, fluorescent protein: $P_A$), the expression of reporter mKate2 (mRNA: $mRNA_B$, immature/ non-fluorescent protein: $PI_B$, fluorescent protein: $P_B$) and the expression of reporter GFP (mRNA: $mRNA_C$, immature/ non-fluorescent protein: $PI_C$, fluorescent protein: $P_C$).

ODE system

sgRNA dynamics

$$\frac{d[sgRNA_1]}{dt} = f_{\left(Ara_{sgRNA_1}\right)} + b_{sgRNA_1} - d_{RNA} \cdot \left[sgRNA_1\right] - k_{f_{ds}} \cdot \left[dCas\right] \cdot \left[sgRNA_1\right] + k_{r_{ds}} \cdot [dCas : sgRNA_1]$$

$$\frac{d[sgRNA_2]}{dt} = f_{\left(Ara_{sgRNA2}\right)} + b_{sgRNA_2} - d_{RNA} \cdot \left[sgRNA_2\right] - k_{f_{ds}} \cdot \left[dCas\right] \cdot \left[sgRNA_2\right] + k_{r_{ds}} \cdot [dCas : sgRNA_2]$$

$$\frac{d[dCas : sgRNA_1]}{dt} = k_{f_{ds}} \cdot \left[dCas\right] \cdot \left[sgRNA_1\right] + k_{r_{dsd1}} \cdot \left[dCas : sgRNA_1 : DNA_C\right]$$
$$-m \cdot \left[dCas : sgRNA_1\right] - k_{r_{ds}} \cdot \left[dCas : sgRNA_1\right] - k_{f_{dsd1}} \cdot \left[dCas : sgRNA_1\right] \cdot \left[DNA_C\right]$$

$$\frac{d[dCas : sgRNA_2]}{dt} = k_{f_{ds}} \cdot \left[dCas\right] \cdot \left[sgRNA_2\right] + k_{r_{dsd2}} \cdot \left[dCas : sgRNA_2 : DNA_B\right]$$
$$-m \cdot \left[dCas : sgRNA_2\right] - k_{r_{ds}} \cdot \left[dCas : sgRNA_2\right] - k_{f_{dsd2}} \cdot \left[dCas : sgRNA_2\right] \cdot \left[DNA_B\right]$$

$$\frac{d\left[dCas : sgRNA_1 : DNA_C\right]}{dt} = k_{f_{dsd1}} \cdot \left[dCas : sgRNA_1\right] \cdot \left[DNA_C\right] - k_{r_{dsd1}} \cdot \left[dCas : sgRNA_1 : DNA_C\right]$$
$$-m \cdot \left[dCas : sgRNA_1 : DNA_C\right]$$

$$\frac{d\left[dCas : sgRNA_2 : DNA_B\right]}{dt} = k_{f_{dsd2}} \cdot \left[dCas : sgRNA_2\right] \cdot \left[DNA_B\right] - k_{r_{dsd2}} \cdot \left[dCas : sgRNA_2 : DNA_B\right]$$
$$-m \cdot \left[dCas : sgRNA_2 : DNA_B\right]$$

$$\frac{d[sgRNA_3]}{dt} = k_{sgRNA_3} \cdot \left[DNA_B\right] - d_{RNA} \cdot \left[sgRNA_3\right] - \mathrm{k}_{\mathrm{f}_{\mathrm{ds}}} \cdot \left[\mathrm{dCas}\right] \cdot \left[\mathrm{sgRNA}_3\right] + \mathrm{k}_{\mathrm{r}_{\mathrm{ds}}} \cdot [\mathrm{dCas} : \mathrm{sgRNA}_3]$$

$$\frac{d[dCas : sgRNA_3]}{dt} = k_{f_{ds}} \cdot \left[dCas\right] \cdot \left[sgRNA_3\right] + k_{r_{dsd3}} \cdot \left[dCas : sgRNA_3 : DNA_C\right]$$
$$-m \cdot \left[dCas : sgRNA_3\right] - k_{r_{ds}} \cdot \left[dCas : sgRNA_3\right] - k_{f_{dsd3}} \cdot \left[dCas : sgRNA_3 \cdot DNA_C\right]$$

$$\frac{d\left[dCas : sgRNA_3 : DNA_C\right]}{dt} = k_{f_{dsd3}} \cdot \left[dCas : sgRNA_3\right] \cdot \left[DNA_C\right]$$
$$- k_{r_{dsd3}} \cdot \left[dCas : sgRNA_3 : DNA_C\right] - m \cdot \left[dCas : sgRNA_3 : DNA_C\right]$$

Reporter protein dynamics

$$\frac{d[mRNA_A]}{dt} = f_{\left(Ara_{mRNA_A}\right)} + b_{mRNA_A} - d_{RNA} \cdot [mRNA_A]$$

$$\frac{d[PI_A]}{dt} = k_{PI_A} \cdot [mRNA_A] - m_{PI_A} \cdot [PI_A] - (m + d_{PI_A}) \cdot [PI_A]$$

$$\frac{d[P_A]}{dt} = m_{PI_A} \cdot [PI_A] - (m + d_{PI_A}) \cdot [P_A]$$

$$\frac{d[mRNA_B]}{dt} = k_{mRNA_B} \cdot [DNA_B] - d_{RNA} \cdot [mRNA_B]$$

$$\frac{d[PI_B]}{dt} = k_{PI_B} \cdot [mRNA_B] - m_{PI_B} \cdot [PI_B] - (m + d_{PI_B}) \cdot [PI_B]$$

$$\frac{d[P_B]}{dt} = m_{PI_B} \cdot [PI_B] - (m + d_{PI_B}) \cdot [P_B]$$

$$\frac{d[mRNA_C]}{dt} = k_{mRNA_C} \cdot [DNA_C] - d_{RNA} \cdot [mRNA_C]$$

$$\frac{d[PI_C]}{dt} = k_{PI_C} \cdot [mRNA_C] - m_{PI_C} \cdot [PI_C] - (m + d_{PI_C}) \cdot [PI_C]$$

$$\frac{d[P_C]}{dt} = m_{PI_C} \cdot [PI_C] - (m + d_{PI_C}) \cdot [P_C]$$

Algebraic equations

$$[dCas] = [dCas_{total}] - [dCas:sgRNA_1] - [dCas:sgRNA_1:DNA_C] - [dCas:sgRNA_2]$$
$$- [dCas:sgRNA_2:DNA_B] - [dCas:sgRNA_3] - [dCas:sgRNA_3:DNA_C]$$

$$[DNA_B] = [DNA_{B_{total}}] - [dCas:sgRNA_2:DNA_B]$$

$$[DNA_C] = [DNA_{C_{total}}] - [dCas:sgRNA_1:DNA_C] - [dCas:sgRNA_3:DNA_C]$$

$$\mathrm{f}_{(\mathrm{Ara}_{\mathrm{sgRNA}_1})} = \frac{Ara^n}{Ara^n + K_m{}^n} \cdot k_2 \cdot [\mathrm{DNA}_{A_{total}}] \cdot (1 - \mathrm{b}_{\mathrm{sgRNA}_1})$$

$$\mathrm{f}_{(\mathrm{Ara}_{\mathrm{sgRNA}_2})} = \frac{Ara^n}{Ara^n + K_m{}^n} \cdot k_2 \cdot [\mathrm{DNA}_{B_{total}}] \cdot (1 - \mathrm{b}_{\mathrm{sgRNA}_2})$$

$$\mathrm{f}_{(\mathrm{Ara}_{\mathrm{mRNA}_\mathrm{A}})} = \frac{Ara^n}{Ara^n + K_m{}^n} \cdot k_2 \cdot [\mathrm{DNA}_{A_{total}}] \cdot (1 - \mathrm{b}_{\mathrm{mRNA}_\mathrm{A}})$$

$$G(t) = p_1 + \sum_{k=1}^{3} \frac{p_2^k}{\left(1 + p_3^k e^{-t \cdot p_4^k}\right)^{1/p_5^k}}$$

$$m(t) = \frac{dG(t)}{dt}$$

A 3-node topology model with three inhibitions including a mutual inhibition

Here we present the ODE system with the algebraic functions for the 3-node model, assuming that we have:

- Three inhibitions via expression of $sgRNA_1$, $sgRNA_2$ and $sgRNA_3$ and the corresponding dCas complexes. Arabinose induces the expression of $sgRNA_1$ that inhibits the expression of the constitutively expressed green node. In addition, the green node is inhibited by $sgRNA_2$, which is expressed by the blue node. Finally, the green node inhibits the expression of the orange node by expressing $sgRNA_3$.
- The expression of reporter mKO2 (mRNA: $mRNA_A$, immature/ non-fluorescent protein: $PI_A$, fluorescent protein: $P_A$), the expression of reporter mKate2 (mRNA: $mRNA_B$, immature/ non-fluorescent protein: $PI_B$, fluorescent protein: $P_B$) and the expression of reporter GFP (mRNA: $mRNA_C$, immature/ non-fluorescent protein: $PI_C$, fluorescent protein: $P_C$).

ODE system

sgRNA dynamics

$$\frac{d[sgRNA_1]}{dt} = f_{\left(Ara_{sgRNA_1}\right)} + b_{sgRNA_1} - d_{RNA}\cdot[sgRNA_1] - k_{f_{ds}}\cdot[dCas]\cdot[sgRNA_1] + k_{r_{ds}}\cdot[dCas:sgRNA_1]$$

$$\frac{d[sgRNA_2]}{dt} = k_{sgRNA_2}\cdot[DNA_B] - d_{RNA}\cdot[sgRNA_2] - \mathrm{k}_{\mathrm{f}_{\mathrm{ds}}}\cdot[\mathrm{dCas}]\cdot[\mathrm{sgRNA}_2] + \mathrm{k}_{\mathrm{r}_{\mathrm{ds}}}\cdot[\mathrm{dCas:sgRNA}_2]$$

$$\frac{d[dCas:sgRNA_1]}{dt} = k_{f_{ds}}\cdot[dCas]\cdot[sgRNA_1] + k_{r_{dsd1}}\cdot[dCas:sgRNA_1:DNA_C]$$
$$-m\cdot[dCas:sgRNA_1] - k_{r_{ds}}\cdot[dCas:sgRNA_1] - k_{f_{dsd1}}\cdot[dCas:sgRNA_1]\cdot[DNA_C]$$

$$\frac{d[dCas:sgRNA_2]}{dt} = k_{f_{ds}}\cdot[dCas]\cdot[sgRNA_2] + k_{r_{dsd2}}\cdot[dCas:sgRNA_2:DNA_C]$$
$$-m\cdot[dCas:sgRNA_2] - k_{r_{ds}}\cdot[dCas:sgRNA_2] - k_{f_{dsd2}}\cdot[dCas:sgRNA_2]\cdot[DNA_C]$$

$$\frac{d[dCas:sgRNA_1:DNA_C]}{dt} = k_{f_{dsd1}}\cdot[dCas:sgRNA_1]\cdot[DNA_C] - k_{r_{dsd1}}\cdot[dCas:sgRNA_1:DNA_C]$$
$$-m\cdot[dCas:sgRNA_1:DNA_C]$$

$$\frac{d\left[dCas:sgRNA_2:DNA_C\right]}{dt}=k_{f_{dsd2}}\cdot\left[dCas:sgRNA_2\right]\cdot\left[DNA_C\right]-k_{r_{dsd2}}\cdot\left[dCas:sgRNA_2:DNA_C\right]$$
$$-m\cdot\left[dCas:sgRNA_2:DNA_C\right]$$

$$\frac{d[sgRNA_3]}{dt}=k_{sgRNA_3}\cdot\left[DNA_C\right]-d_{RNA}\cdot\left[sgRNA_3\right]-\mathrm{k}_{\mathrm{f}_{\mathrm{ds}}}\cdot\left[\mathrm{dCas}\right]\cdot\left[\mathrm{sgRNA}_3\right]+\mathrm{k}_{\mathrm{r}_{\mathrm{ds}}}\cdot[\mathrm{dCas}:\mathrm{sgRNA}_3]$$

$$\frac{d[dCas:sgRNA_3]}{dt}=k_{f_{ds}}\cdot\left[dCas\right]\cdot\left[sgRNA_3\right]+k_{r_{dsd3}}\cdot\left[dCas:sgRNA_3:DNA_A\right]$$
$$-m\cdot\left[dCas:sgRNA_3\right]-k_{r_{ds}}\cdot\left[dCas:sgRNA_3\right]-k_{f_{dsd3}}\cdot\left[dCas:sgRNA_3\cdot DNA_A\right]$$

$$\frac{d\left[dCas:sgRNA_3:DNA_A\right]}{dt}=k_{f_{dsd3}}\cdot\left[dCas:sgRNA_3\right]\cdot\left[DNA_A\right]$$
$$-\ k_{r_{dsd3}}\cdot\left[dCas:sgRNA_3:DNA_A\right]-m\cdot\left[dCas:sgRNA_3:DNA_A\right]$$

Reporter protein dynamics

$$\frac{d[mRNA_A]}{dt}=f_{\left(Ara_{mRNA_A}\right)}+b_{mRNA_A}-d_{RNA}\cdot[mRNA_A]$$

$$\frac{d[PI_A]}{dt}=k_{PI_A}\cdot\left[mRNA_A\right]-m_{PI_A}\cdot\left[PI_A\right]-\left(m+d_{PI_A}\right)\cdot\left[PI_A\right]$$

$$\frac{d[P_A]}{dt}=m_{PI_A}\cdot\left[PI_A\right]-\left(m+d_{PI_A}\right)\cdot\left[P_A\right]$$

$$\frac{d[mRNA_B]}{dt}=k_{mRNA_B}\cdot\left[DNA_B\right]-d_{RNA}\cdot[mRNA_B]$$

$$\frac{d[PI_B]}{dt}=k_{PI_B}\cdot\left[mRNA_B\right]-m_{PI_B}\cdot\left[PI_B\right]-\left(m+d_{PI_B}\right)\cdot\left[PI_B\right]$$

$$\frac{d[P_B]}{dt}=m_{PI_B}\cdot\left[PI_B\right]-\left(m+d_{PI_B}\right)\cdot\left[P_B\right]$$

$$\frac{d[mRNA_C]}{dt}=k_{mRNA_C}\cdot\left[DNA_C\right]-d_{RNA}\cdot[mRNA_C]$$

$$\frac{d[PI_C]}{dt}=k_{PI_C}\cdot\left[mRNA_C\right]-m_{PI_C}\cdot\left[PI_C\right]-\left(m+d_{PI_C}\right)\cdot\left[PI_C\right]$$

$$\frac{d[P_C]}{dt}=m_{PI_C}\cdot\left[PI_C\right]-\left(m+d_{PI_C}\right)\cdot\left[P_C\right]$$

Algebraic equations

$$\left[dCas\right]=\left[dCas_{total}\right]-\left[dCas:sgRNA_1\right]-\left[dCas:sgRNA_1:DNA_C\right]-\left[dCas:sgRNA_2\right]$$
$$-\left[dCas:sgRNA_2:DNA_C\right]-\left[dCas:sgRNA_3\right]-\left[dCas:sgRNA_3:DNA_A\right]$$

$$\left[DNA_A\right]=\left[DNA_{A_{total}}\right]-\left[dCas:sgRNA_3:DNA_A\right]$$

$$\left[DNA_C\right]=\left[DNA_{C_{total}}\right]-\left[dCas:sgRNA_1:DNA_C\right]-\left[dCas:sgRNA_2:DNA_C\right]$$

$$\mathrm{f}_{\left(\mathrm{Ara}_{\mathrm{sgRNA}_1}\right)}=\frac{Ara^n}{Ara^n+K_m{}^n}\cdot k_2\cdot[\mathrm{DNA}_{A_{total}}]\cdot\left(1-\mathrm{b}_{\mathrm{sgRNA}_1}\right)$$

$$\mathrm{f}_{\left(\mathrm{Ara}_{\mathrm{mRNA}_\mathrm{A}}\right)}=\frac{Ara^n}{Ara^n+K_m{}^n}\cdot k_2\cdot[\mathrm{DNA}_{A_{total}}]\cdot\left(1-\mathrm{b}_{\mathrm{mRNA}_\mathrm{A}}\right)$$

$$G\left(t\right)=p_1+\sum_{k=1}^{3}\frac{p_2^k}{\left(1+p_3^k e^{-t\cdot p_4^k}\right)^{1/p_5^k}}$$

$$m\left(t\right)=\frac{dG\left(t\right)}{dt}$$
